# Concentration- and time-dependence of phosphorylation events stimulated by fibroblast growth factor-4 (FGF-4) in Rama 27 fibroblasts

**DOI:** 10.1101/2025.05.09.652961

**Authors:** Marim M. Alsadun, Dunhao Su, Edwin A. Yates, David G. Fernig

## Abstract

Gradients of FGFs are an essential feature of many developmental processes. Thus, FGF2 stimulates different responses at high and low concentrations, while FGF4 gradients are critical, e.g., in limb development., but it is not known if responses to FGF4 differ in a concentration-dependent manner. Employing rat mammary (Rama) 27 fibroblasts, we therefore measured the FGF4 concentration- and time-dependence of the simulation of phosphorylation of FRS2α, MAPK1 and MAPK3 which are downstream of the FGF receptor (FGFR1c). At 10 pg/mL FGF4 caused a very weak phosphorylation of FRS2α at Y435 and Y196 and a stronger one of MAPK1 and MAPK3 that oscillated with maximum levels after 30 min and 240 min stimulation. The phosphorylation of these proteins at 1 ng/mL FGF4 was considerably stronger and showed a similar oscillation. At 10 ng/mL FGF4, the phosphorylation of FRS2α reached an early peak after 5 min, declined at 15 min and then rose again at 30 min before declining to the end of the time-course at 240 min, whereas the phosphorylation of MAPK1 and MAPK3 was strong after 5 minutes, reached a maximum after 30 minutes and then declined gradually. At 1 µg/mL and 3 µg/mL FGF4 after 15 min and 60 min the phosphorylation of FRS2α, MAPK1 and MAPK3 was much lower at 60 min compared to that observed at lower concentrations of FGF4. The data indicate that at low concentrations FGF4 elicits an oscillatory response at the level of phosphorylation of FRS2α and of MAPK1 and MAPK3, whereas a bell-shaped dose response may occur at the highest concentrations of FGF4. The addition of exogenous heparin, an effective mimic of endogenous heparan sulfate, suggests that there may be an influence of the interaction of FGF4 with pericellular matrix and the dose-and time-dependence of the phosphorylation of FRS2α and MAPK1and MAPK3.

## Introduction

Fibroblast Growth Factors (FGFs) are widely expressed in vertebrates and invertebrates and regulate fundamental functions in development and homeostasis in relation to morphogenesis, growth and maintenance of organs and body structures[1]. Unsurprisingly, signalling activated by FGF ligands also contributes to the pathology of many non-communicable diseases such as cancers[2], whereas the FGF ligands themselves possess considerable potential a tissue repair therapeutics[3] The paracrine FGF ligands elicit intracellular signals by forming a ternary complex with their glycosaminoglycan heparan sulfate (HS) co-receptor and their receptor tyrosine kinases, FGFR[1] [2, 4-7]. The engagement of FGF ligand and HS co-receptor leads to dimerization of the FGFR and activation of its intracellular tyrosine kinase. Subsequently, phosphorylation of tyrosine residues in the cytosolic part of the receptor and on the adaptor protein fibroblast growth factor receptor substrate 2α (FRS2α) provide docking sites for proteins that activate downstream signalling pathways such as the GRB2-MAPK and PI3K-AKT axes [1, 2, 8]. Thus, once phosphorylated, FRS2α acts as a further platform for docking SH2 domain-containing proteins, including GRB2 and SHP-2, which are crucial for downstream signalling processes[7-9]. Specifically, Tyr436 is essential for the efficient recruitment of SHP-2, while Tyr196 serves as a docking site for GRB2-SOS complexes.

FGF4 is a paracrine FGF4, which in common with other members of this subfamily binds the IIIc splice variants of FGFR1 to -3, with a marked preference for FGFR1c and FGFR2c over FGFR3c or FGFRΔ4 if the concentration required for half maximal stimulation of the Baf3 cells is considered [10]. This contrasts with FGF2, which has a preference for FGFR1c and FGFR3c over FGFR2c. Upon FGFR activation, FRS2 undergoes tyrosine phosphorylation[6-8, 11]. FGF4 is essential for early mouse embryonic development, regulating cell division in both embryonic and extraembryonic tissues [12, 13]. Its absence leads to embryonic lethality shortly after implantation, highlighting its critical role in early embryogenesis[12, 13]. Thereafter, FGF4 plays a critical role in diversifying valve precursor cells into cartilage and tendon cell lineages during heart valve development in chick embryos[14] and is an essential signalling component of the apical ectodermal ridge where it contributes to the outgrowth of the limb bud [15].

Intracellular signals stimulated by FGF-4 have been analysed in a variety of cell types. In human embryonic stem cell (ESC) cultures FGF4 at 10 ng/mL stimulated sustained phosphorylation of mitogen-activated protein kinases-1 and -3 (MAPK-1, -3, also known as extracellular regulated kinases ERK-1 and -2) over 24 h. In contrast, in pituitary cells where FGF4 stimulated prolactin transcription through a Rac1-dependent mechanism, involving PLCγ signalling, independent of the Ras-Raf pathway[16, 17] FGF-4 stimulated transient phosphorylation of MAPK-1 and -3 [17]. More commonly a single concentration of FGF4 has been used, for example 10 ng/ml in [13], 50 ng/mL in experiments involving O-GlcNAc transferase knockdown cells [13] and 1 µg/mL in experiments with bovine embryos [18]. In experiments utilizing NIH3T3 fibroblasts as a model system [19], it was discovered that the most effective concentration for promoting cell division was approximately 1 ng/ml of FGF4, with up to 50 ng/mL having slightly lower efficacy [19]. There are, however, no detailed analyses of the dose- or time-dependence of intracellular signals elicited by FGF4.

In contrast, the dose- and time-dependence of cell responses to FGF1 and FGF2 have been examined in detail. Thus, it is well established that the simulation of cell division by FGF1 and FGF2 has a bell-shaped dose response in cultured cells [20-22]. In the case of FGF2 in fibroblasts this has been demonstrated to arise from high concentrations of FGF2 failing to cause stimulation of the phosphorylation of the adaptor FRS2α the consequence of which is that the stimulation of the phosphorylation of MAPK-1 and -3 is transient, which is insufficient to stimulate cell division [9, 11]. The distinct responses of cells to lower and higher concentrations of FGF2 are likely to explain how high and low concentrations of FGFs are able to pattern the ventral foregut into liver and lung [23]. Since a focal source of FGF4 is required for limb outgrowth[15, 24] it was of interest to determine whether the stimulation of phosphorylation of FRS2α, MAPK1 and MAPK3 by FGF4, like that by FGF1 and FGF2 elicited a bell-shaped dose response.

We therefore used Rat mammary (Rama) 27 fibroblasts as a model system to interrogate the dose-and time-dependence of the stimulation of phosphorylation of FRS2α, MAPK-1 and -3 and AKT to enable a direct comparison with previous work with FGF2 on these cells [9]. The data demonstrate that FGF4 stimulate phosphorylation of FRS2α, MAPK-1 and -3 and AKT. Moreover, the kinetics of phosphorylation of these proteins is oscillatory at lower concentration of FGF4. At concentrations of 1 µg/mL FGF4 and above the level of phosphorylation observed after 60 min is attenuated, whereas that at 15 min is largely unaffected. The data suggest that FGF4 may elicit concentration- and time-dependent downstream phosphorylation of FRS2α, MAPK-1 and -3 and AKT, which may provide distinct cellular responses to a gradient of FGF4.

## Materials and Methods

### Materials

FGF4 was obtained from Dr.Yong Li (University of Liverpool, UK), Cell culture reagents were Dulbecco’s modified Eagle’s medium (DMEM, Gibco Life Technologies), foetal bovine serum (FBS),(Gibco), L-glucosamine (Gibco), insulin (Sigma-Aldrich), hydrocortisone (Sigma-Approach), bovine serum, albumin (BSA; Sigma-Aldrich catalogue number A7030), phosphate-buffered saline (PBS; 137 mM NaCl, 2.7 mM KCl, 8 mM Na_2_HPO_4_, 1.5 mM KH_2_PO_4_, pH 7.2, 25 mL Versene (Gibco) containing 2.5 % (w/v) trypsin, 100 mm diameter cell culture dishes (cyto one, Roskilde, Denmark), protease inhibitor cocktail, 1 mM sodium orthovanadate, and phosphatase inhibitors were from Sigma-Aldrich. Reagents for SDS-PAGE and electro transfer were obtained from multiple sources. Acrylamide/bis-acrylamide solution (30% w/v, 29:1) was purchased from Seven Biotech Ltd (Cat. No. 20-9001-10) and a nitrocellulose membrane (0.45 µm, Bio-Rad, Cat# 1620145). Other components, including the resolving gel buffer, stacking gel buffer, 10% SDS stock, running buffer, and transfer buffer, were prepared in-house according to standard protocols, antibody to phosphotyrosine phospho-FRS2-α (Tyr196) antibody #3864, Phospho-FRS2-α (Tyr436) antibody #3861, and to phospho-MAPK1 and 3 antibody #9102 and to GAPDH (D16H11) rabbit mAb #5174 were from Cell Signalling Technology (Hitchin, UK). Secondary peroxidase-labelled anti-IgG antibodies (anti-rabbit and anti-mouse, Antibody #7074, and Antibody #7076, respectively were from Cell Signalling Technology (Hitchin, UK).

### Stimulation of phosphorylation of MAPK1, MAPK3 and FRS2α by FGF4

Rama (rat mammary) 27 fibroblast cells [25] were plated in sterile 100 mm diameter cell culture dishes and incubated at 37 °C with 10 ml Routine medium (RM, DMEM (Dulbecco’s modified Eagle’s medium supplemented with 10 % (v/v) FBS (foetal bovine serum) (Gibco), 20 mM L-glucosamine, 50ng/ml insulin, 50 ng/mL hydrocortisone. Rama 27 cells at 70% confluency were washed twice with PBS and then 10 mL Step Down Medium (SDM, DMEM supplemented with 250 µg/ml BSA) was added. After 20-24 hours, 9 mL fresh SDM was added and cells incubated in this for an additional 3-4 hours. FGF4 was then added in 1 mL SDM at the concentrations and for the times indicted in the figure legends. Cells were washed twice with ice-cold Tris-buffered saline (TBS, pH 7.2) containing 0.1 mM sodium orthovanadate, and then scraped (on ice) into 200 µL SDS-PAGE sample buffer (without bromophenol blue) supplemented with 1 mM sodium orthovanadate, 1 tablet of protease inhibitor cocktail and 1 tablet of phosphatase inhibitor cocktail per 10 mL. The cell lysate was then sonicated for 10 s and heated for 5 min at 99 °C.

### SDS-PAGE and Western Blotting

#### Phosphorylation signalling detection for FRS2-α and phospho-p44/42 MAPK (Erk1/2) (T202/Y204)

Identical amounts of protein were loaded onto a 10 % (w/v) polyacrylamide gel. Polypeptides were separated by electrophoresis and then transferred to a nitrocellulose membrane (0.45 µm, Bio-Rad, Cat# 1620145). Transfer buffer was prepared by diluting 10X transfer buffer stock (containing 250 mM Tris base PH 8.6, 1.92 M glycine, and 1% (w/v) SDS) 1:10 with distilled water, followed by the addition of 15% (v/v) methanol to the final volume. The membrane was incubated in Ponceau S red staining solution (#102582210, Merck) for 2 minutes, then washed twice with TBS supplemented with 0.1% (v/v) Tween-20 (TBST) for 10 minutes. This was followed by blocking with 5% (w/v) non-fat dried milk in TBST for at least 2 hours, and two additional washes with TBST (5 minutes each). The membrane was then incubated overnight at 4 °C with rabbit polyclonal antibodies to FRS2α phosphorylated on Tyr436 and on Tyr196 (1:5,000, in 10 mL TBST containing 2% (w/v) BSA). After two 5-minute TBST washes, membranes were incubated for 180 minutes at room temperature with HRP-conjugated anti-rabbit IgG secondary antibody (1:5,000; #7074, Cell Signalling Technology, UK), prepared in TBST containing 2% BSA. For detection of dual-phosphorylated MAPK1/3 (Thr202/Tyr204), membranes were incubated overnight at 4 °C with a mouse monoclonal antibody (1:10,000; #9102, Cell Signalling Technology) diluted in TBST containing 2% (w/v) BSA. After washing with TBST (3 × 5 minutes), membranes were incubated with HRP-conjugated anti-mouse IgG (1:20,000; #7076S, CST, UK) for 90 minutes at room temperature.

For assessment of protein loading, membranes were reprobed with rabbit monoclonal anti-GAPDH antibody (D16H11; 1:10,000; #5174, Cell Signalling Technology, Hitchin, UK), incubated overnight at 4 °C in TBST containing 2% BSA. The membrane was then incubated with HRP-conjugated anti-rabbit IgG (1:10,000; #7074, CST, UK) for 90 minutes at room temperature. HRP activity was visualised using an ECL enhanced chemiluminescence kit (Millipore, #WBKLS0500) according to the manufacturer’s instructions.

## Results

Rama 27 fibroblasts had previously been shown to express mRNA encoding FGFR1[26] [27] and as mesenchymal cells this will be the ‘c’ isoform, which is consistent with the profile of FGF ligands that can stimulate signalling downstream of the FGFR [10]. As expected, western blots of Rama 27 cells demonstrate that they express FGFR1 (Fig. 1). Importantly, no other FGFR was detectable by western blot (Fig. S1). This is a key observation, since while FGF4 and FGF2 have a similar preference for FGFR1, FGF4 has a preference for FGFR2c over FGFR3c, whereas this is reversed for FGF2 [10]. Consequently, downstream signals stimulated by FGF4 will be generated by activation of FGFR1c and any differences noted with those generated by FGF2 are likely to be intrinsic to the FGF ligands, and their interaction with cellular HS, rather than the FGFR.

**Figure 1:**
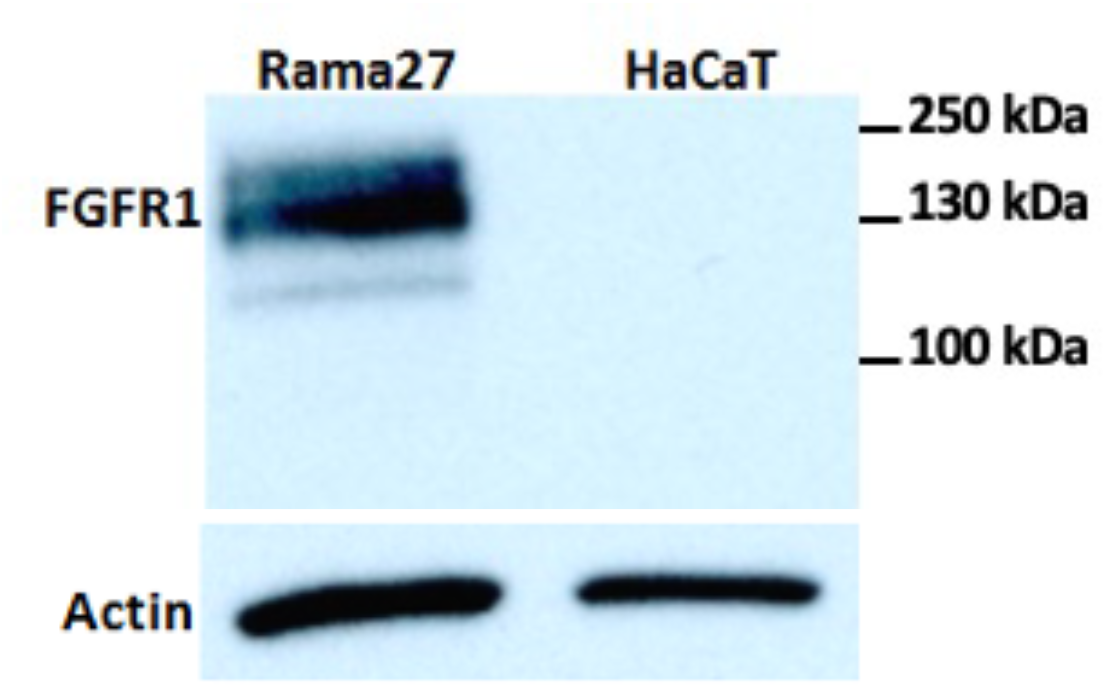
Rama 27 fibroblasts produce FGFR1. Rama 27 fibroblasts were collected by scaping in sample buffer and polypeptides were subjected to SDS-PAGE and then transferred to a nitrocellulose membrane. FGFR1 was detected by western blotting. HaCaT human keratinocytes were used as a negative control.

### Stimulation of intracellular phosphorylation of FRS2α, MAPK1, and MAPK3 by FGF4 in Rama 27 fibroblasts

An initial experiment aimed to establish the dose-dependence of the dual phosphorylation of MAPK1, MAPK3 and of tyrosines on FRS2α. Phosphorylation of these proteins was determined after 15 minutes stimulation of quiescent Rama 27 fibroblasts with a broad range of FGF4 concentrations ranging from 10 pg/mL to 100 ng/mL. Phosphorylation of FRS2α on Y436 was barely detectable at 100 pg/mL FGF4 and then became increasingly strong as the concentration of FGF4 increased (Fig. 2). Phosphorylation of FRS2α was also monitored using an antibody to phosphorylated Y196. This antibody recognises two polypeptides at 92 kDa and 90 kDa (Fig. 2). Given the that the antibody to phosphorylated Y436 (Fig. 2) only detects immunoreactivity when cells were incubated with at least 100 pg/mL FGF4, FRS2α phosphorylated on Y196 would be consistent with the lower immunoreactive band, since this was weakest in the control and stronger when cells were incubated with 10 pg/mL and 100 pg/mL FGF4 (Fig. 2. The origin of the upper band of immunoreactivity is unknown, but since it is strongest in the control, it is most unlikely to correspond to FRS2α that is phosphorylated on Y196, although it is intriguing that it declines somewhat from the level observed in the control to that seen at 1 ng/mL FGF4. As for the antibody recognising phosphoY436, phosphorylation of FRS2α detected by the antibody to phosphoY196 increased strongly in cells incubated with 1 ng/mL FGF4 and then again when cells were incubated with 10 ng/mL FGF4 (Fig. 2). The antibody to FRS2α phosphorylated at Y196 is apparently more sensitive

**Figure 2.**
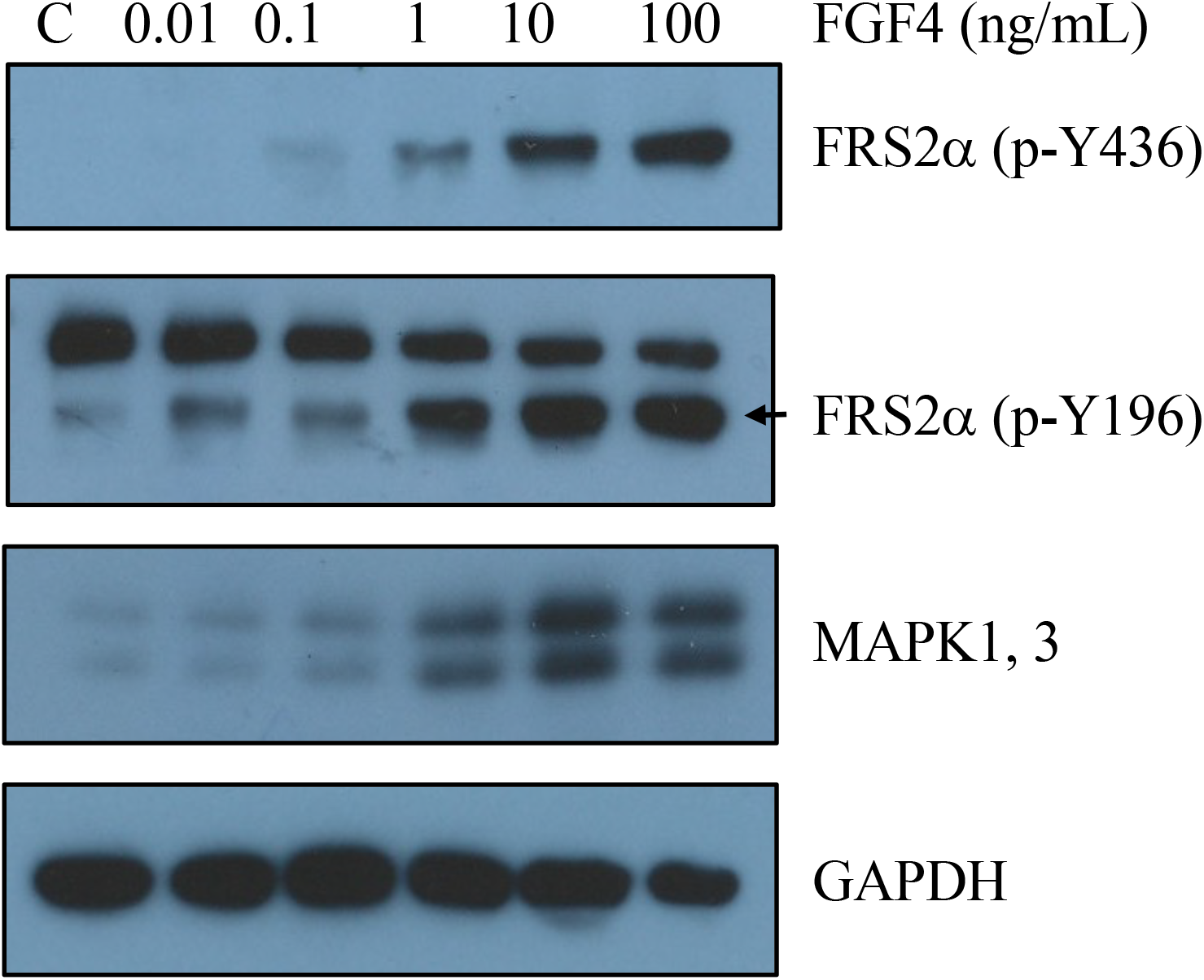
FGF4 stimulates dose-dependent phosphorylation of FRS2α (p-Y436, p-Y196) and MAPK1/3 in Rama 27 fibroblasts. Quiescent Rama 27 fibroblasts were stimulated for 15 minutes with a range of concentrations of FGF4. Cells were scraped into sample buffer, and polypeptides were separated by SDS-PAGE and transferred to a nitrocellulose membrane. Membranes were cut into horizontal strips corresponding to the molecular weights of the target proteins, probed with the appropriate primary and secondary antibodies, and immunoreactivity was detected using enhanced chemiluminescence (ECL).

than that to pY436. However, the former has no off-target immunoreactivity, so consequently both antibodies were used to corroborate any changes observed. The dual phosphorylation of MAPK1 and MAPK3 did not alter appreciably above the levels observed in control cells when cells were stimulated with 10 pg/mL and 100 pg/mL FGF4 but increased substantially with 1 ng/mL FGF4, to reach a maximum at 10 ng/mL FGF4 (Fig. 2).

### Kinetics of phosphorylation ofFRS2α, MAPK1 and -3, and AKT in Rama 27 fibroblasts

Differences in the time-dependence of signalling stimulated by growth factors is an important factor in determining the specificity of the cellular response [28]. In this experiment, quiescent Rama 27 fibroblasts were stimulated over 240 min at low (10 pg/mL, intermediate (1 ng/mL) and high (10 ng/mL) concentrations of FGF4 and the phosphorylation of FRS2α (phosphorylated at Y436 and Y196), MAPK1, MAPK3 and AKT was measured.

#### FRS2α phosphorylation at Y436 and Y196

Consistent with the result obtained previously with 10 pg/mL FGF4 (Fig. 2), at this concentration phosphorylation of FRS2α was undetectable after 15 min stimulation at either Y1436 (Fig. 3A) or Y196 (Fig. 3B). A faint immunoreactive band was observed for phospho Y436 at 30 min and 60 min whereas for phosphoY196 a faint band was detectable at 120 min and 240 min. The level of phosphorylation detected in these instances was near the lower limit of detection. Thus, these data suggest that at 10 pg/mL FGF4 there is at most weak phosphorylation of FRS2α, which may exhibit oscillatory behaviour that differs for the two tyrosines. At the intermediate concentration of FGF4, 1 ng/mL, phosphorylation of FRS2α at Y436 was detectable after 5 min stimulation (Fig. 3C) and remained until 30 min, after which phosphorylation at Y436 declined to undetectable levels, until 240 min, when it became apparent again. The detection of phosphorylation at Y196 followed the same pattern. Strong phosphorylation of FRS2α at Y196 was observed after 5 min stimulation with 1 ng/mL FGF4, and this increased after 15 min. The level of phosphorylation declined slightly after 60 min and reached a minimum at 120 min, before increasing again at 240 min (Fig. 3C). Given the stronger signal with the pY196 antibody, these data suggest that phosphorylation of FRS2α caused by FGF4 has an oscillatory behaviour, which is sufficiently strong to be detected by measurements on a cell population. At 10 ng/ml FGF4, phosphorylation of FRS2α at Y436 was strong after just 5 min stimulation (Fig. 3E), but it then decreased after 15 min, before increasing again at 30 min. Thereafter, there was a shallow decline to the end of the experiment.

**Figure 3:**
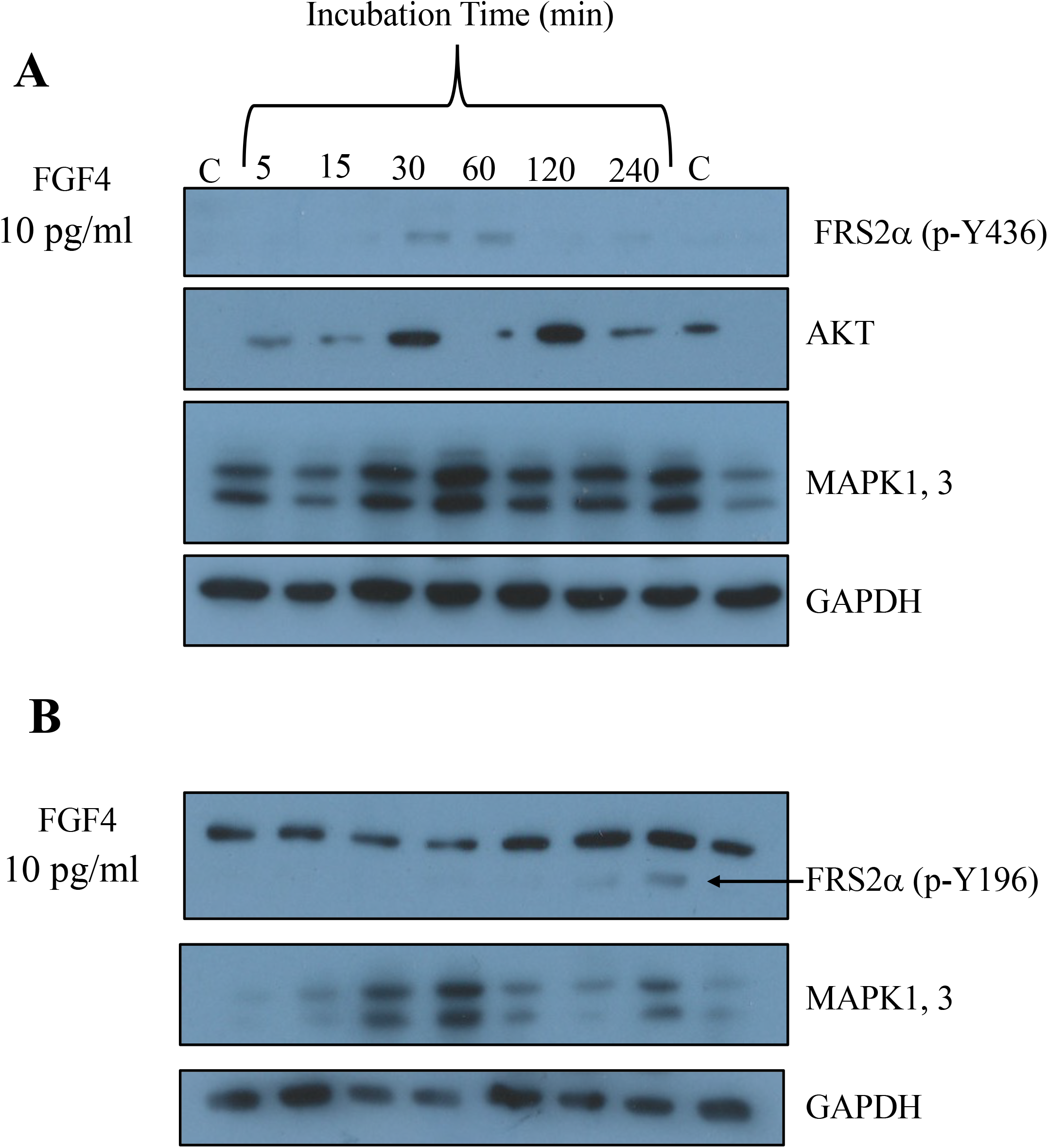
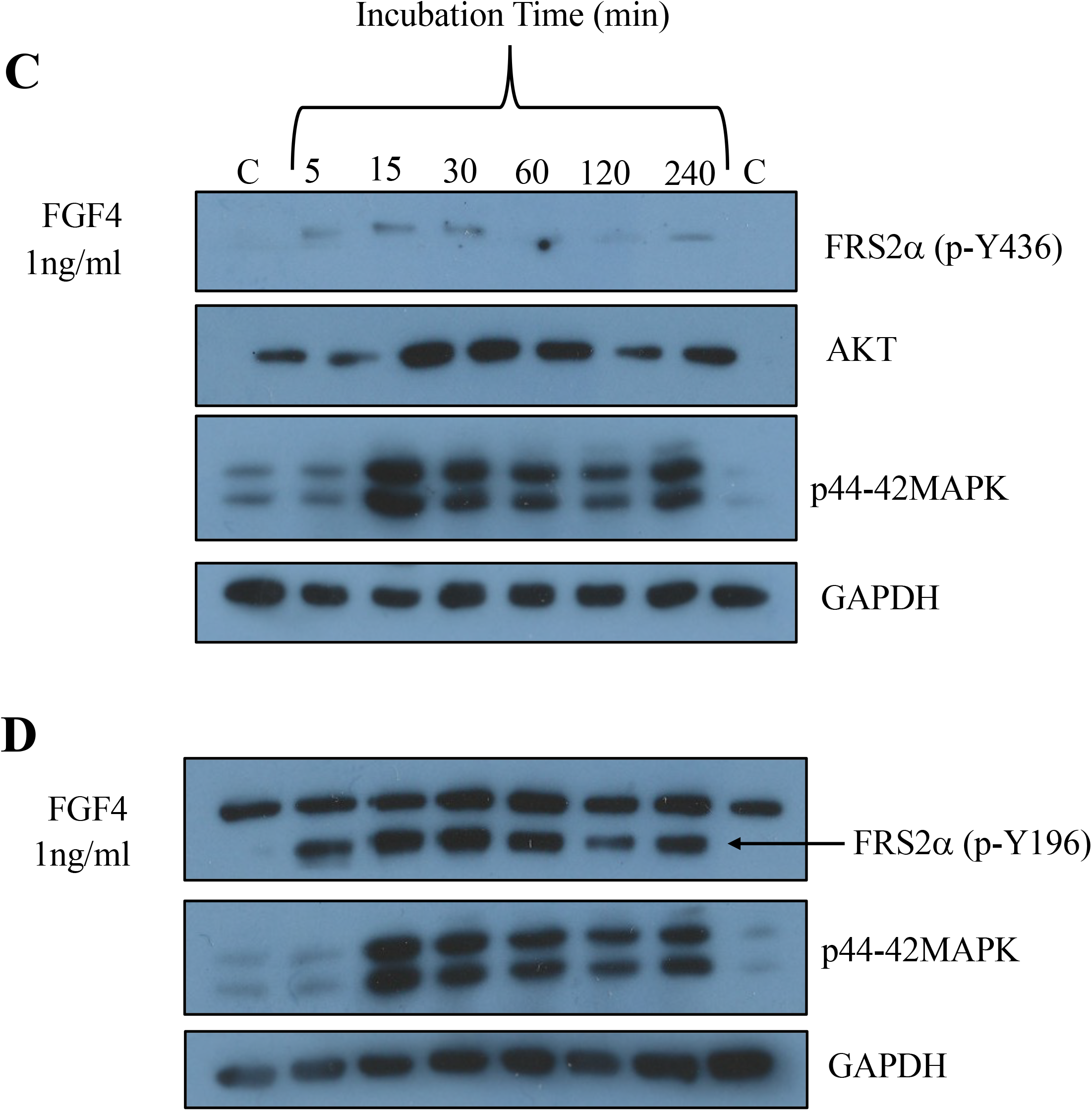
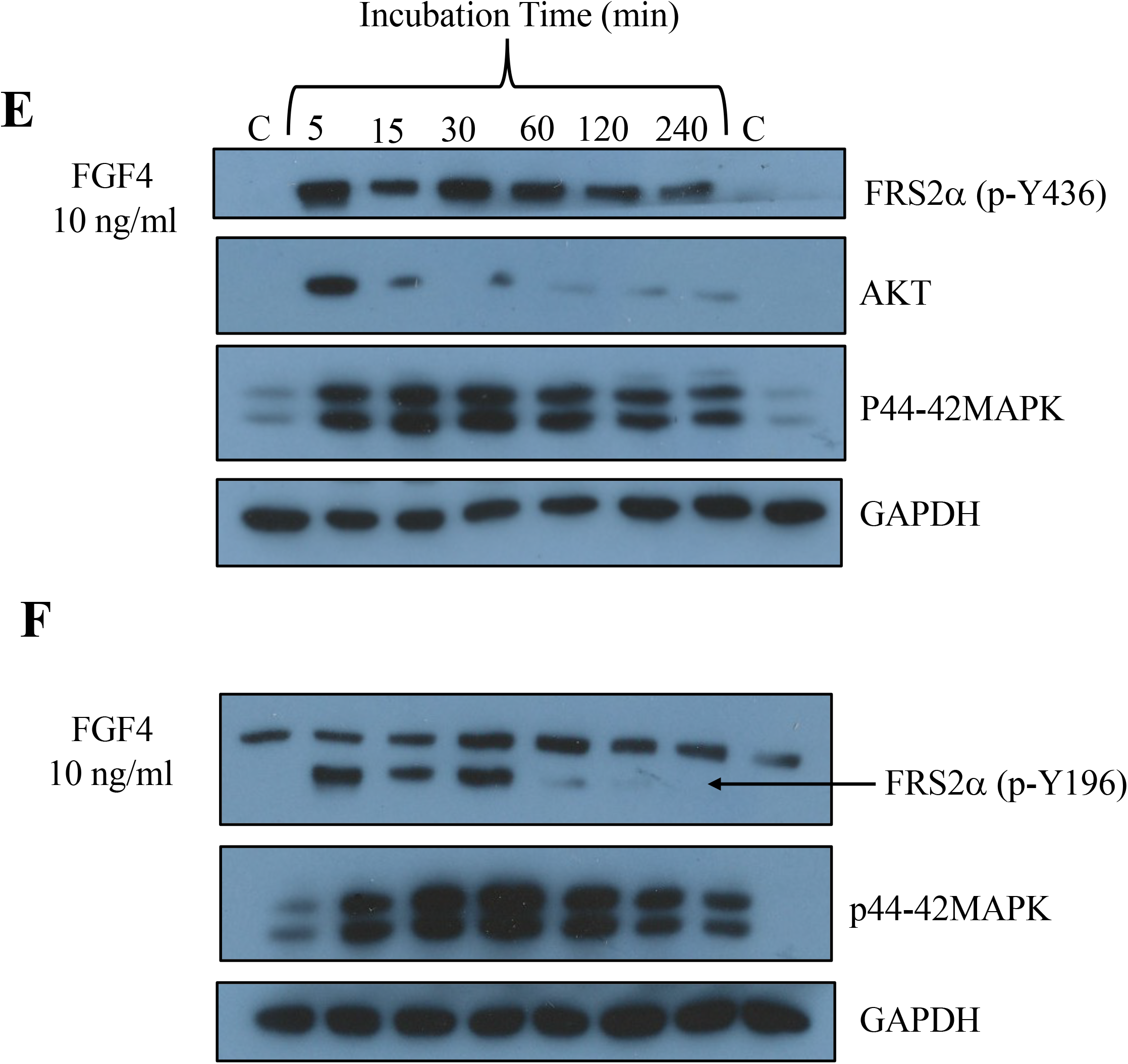
The phosphorylation kinetics of FRS2α (p-Y436, p-Y196), MAPK1 and MAPK3. Rama 27 fibroblasts were stimulated with the indicated concentrations of FGF4 and cells collected by scraping in sample buffer at different times. Following electrophoresis and transfer to a nitrocellulose membrane, the latter was cut into horizontal strips covering the molecular size of the proteins to be probed with the relevant primary and secondary antibodies and immunoreactivity was detected by ECL. Cells were stimulated with (A, B) 10 pg/mL FGF4, (C, D) _ 1 ng/mL FGF4 and (E, F) 10 ng/mL FGF4.

The phosphorylation of FRS2α at Y196 followed the same initial pattern, being strong at 5 min, declining at 15 min and strong again at 30 min (Fig. 3F). However, the level of pY196 FRS2α then declined to undetectable levels from 60 min. These data point to the phosphorylation of FRS2α at Y196 and Y436 oscillating and perhaps having different kinetics of phosphorylation/dephosphorylation.

#### Phosphorylation of MAPK1 and MAPK3

A low but variable basal level of dual phosphorylation of MAPK1 and MAPK3 was observed in control quiescent Rama 27 fibroblasts (Fig. 3). When quiescent Rama 27 fibroblasts were stimulated with 10 pg/mL FGF4 an increase in the dual phosphorylation of MAPK1 and MAPK3 was observed after 15 min, and this increased further by 30 min (Figs 3A, B). By 60 min, the dual phosphorylation of MAPK1 and MAPK3 had declined to near basal levels and remained at this level for another hour (Figs 3A, B). However, after 240 minutes stimulation with 10 pg/ml FGF4, the dual phosphorylation of MAPK1 and MAPK3 increased again (Figs 3A, B). Interestingly, the observed oscillations in the dual phosphorylation of MAPK1 and MAPK3 correlate with the weak phosphorylation of FRS2α at Y196 and Y436 (Figs 3A, B) Stimulation of quiescent Rama 27 fibroblasts with 1 ng/ml FGF4 resulted in a weak stimulation of dual phosphorylation of MAPK1 and MAPK3 after 5 min, and a maximal level of phosphorylation of these proteins after 15 min (Figs 3C, D). The level of dual phosphorylation of MAPK1 and MAPK3 then declined after 30 min to reach a plateau from 60 min to 120 min (Figs 3C, D), after which it increased again at 240 min (Figs 3C, D). There was some correlation with the phosphorylation of FRS2α, since this was lowest at 60 min (Fig. 3C) and 120 min (Figs 3C, D). At the highest concentration of FGF4, 10 ng/ml, dual phosphorylation of MAPK1 and MAPK3 was substantially increased by 5 min and reached a maximum from 15 min to 30 min, after which it declined progressively until 240 min (Figs 3 E, F).

#### AKT phosphorylation

The phosphorylation of AKT represents an important signalling axis, that would be initiated by the phosphorylation of PI3K by the FGFR[8]. Weak basal phosphorylation of AKT was evident in quiescent Rama 27 fibroblasts, which is likely to reflect signalling through default cell survival pathways (Fig. 3). FGF4 at 10 pg/ml stimulated an increase phosphorylation of AKT at 15 min, which then declined at 30 min to basal levels, where it remained at 60 min (Fig. 3A). However, phosphorylation of AKT increased again at 120 min to reach its highest level, and then declined (Fig. 3A). At the higher concentration of 1 ng/ml FGF4 again elicited strong phosphorylation of AKT at 15 min, which declined slightly at 30 min and 60 min, to reach a minimum at 120 min, after which phosphorylation of AKT increased again at 240 min. (Fig. 3C). A very different pattern of phosphorylation of AKT was observed when the cells were stimulated by 10 ng/ml FGF4. Strong phosphorylation of AKT was observed at 5 min, and then by 15 min this had declined markedly to near basal levels, where it remained up to the last time point at 240 min (Fig. 3 E).

### Effects of high concentrations of FGF4 on the phosphorylation of FRS2α, MAPK1 and MAPK3

FGF1 and FGF2 exhibit well documented bell-shaped dose response curves whereby, at high concentrations, which may be attained physiologically in at least some circumstances[15, 23], these FGFs no longer elicit a growth stimulatory response [20] [21, 22], although some aspects of intracellular signalling are retained [7, 9]. At 100 ng/ml FGF4, strong phosphorylation of FRS2α at Y436 and Y196 was detected after 15 min and 60 min, which was maintained at 300 ng/ml (Fig. 4). However, as the concentration of FGF4 increased to 1 µg/ml and then 3 µg/ml, phosphorylation at Y436 declined markedly at both time points. The phosphorylation of FRS2α at Y196 followed a similar pattern, although the relative decline observed at 1 µg/ml and 3 µg/ml was less marked, which may reflect differences in the affinity of the two antibodies (Fig. 4). The dual phosphorylation of MAPK1 and MAPK3 at 15 min was highest at 100 µg/mL FGF4, but only declined slightly at the higher concentrations of the growth factor. At 60 min, the dual phosphorylation of MAPK1 and MAPK3 plateaued at 100 ng/mL to 300 ng/mL, was lower at 1 µg/mL FGF4, and decreased again at 3 µg/mL (Fig. 4). These data indicate that the phosphorylation of FRS2α, MAPK1 and MAPK3 stimulated by FGF4 may have a bell-shaped dose response. However, in contrast to FGF2 in the same cells, not only are considerably higher concentrations of FGF4 required to elicit a decline in phosphorylation of these proteins, but there may be some uncoupling of the amplitude of phosphorylation of FGFR2α and that of MAPK1 and MAPK3.

**Figure 4:**
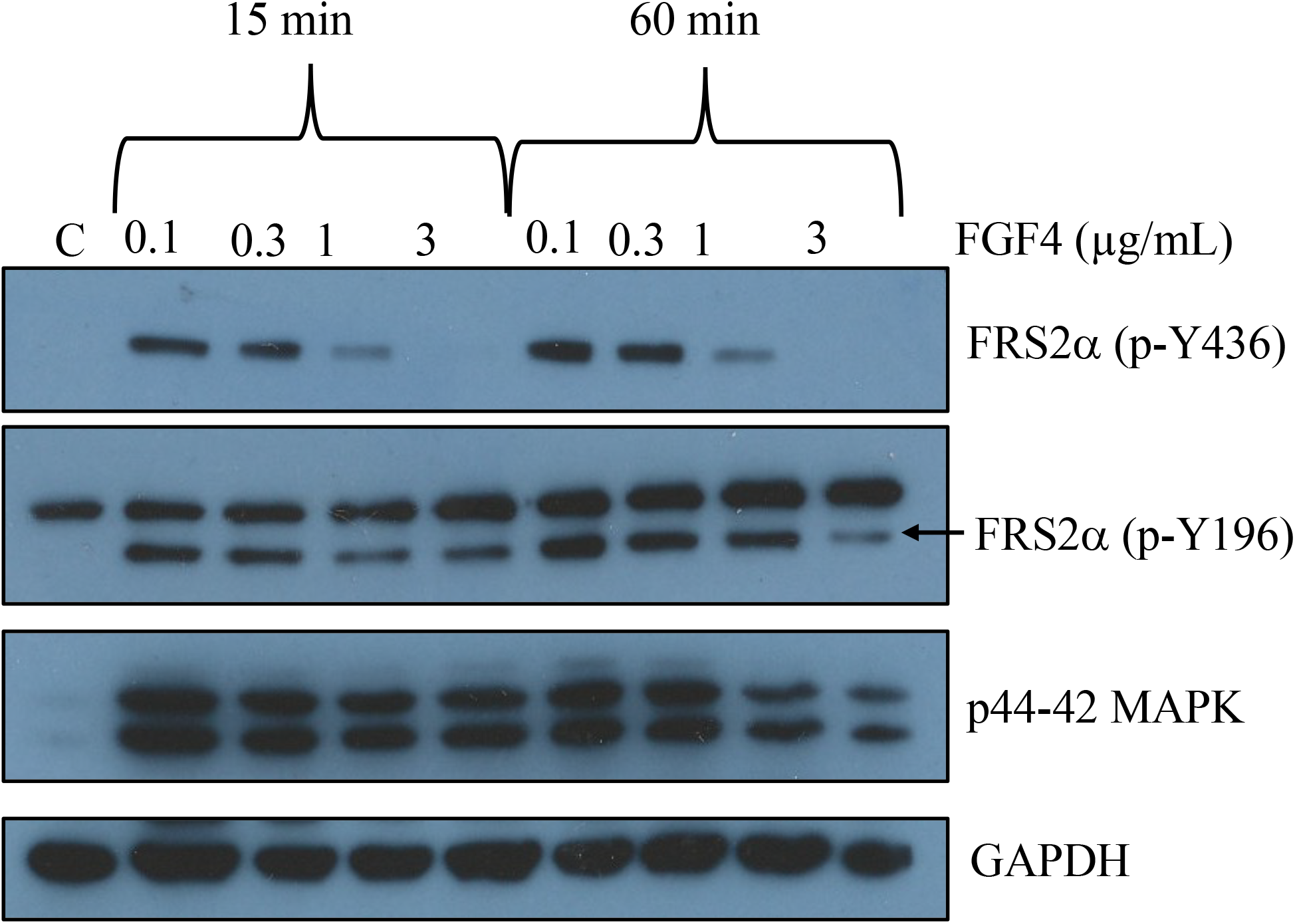
The stimulation of phosphorylation of FRS2α (p-Y436, p-Y196), MAPK1 and MAPK3 at high concentrations of FGF4. Rama 27 fibroblasts were stimulated with the indicated concentrations of FGF4 for 15 min and 60 min and then were cells collected by scraping in sample buffer at different times. After SDS-PAGE and transfer to a nitrocellulose membrane this was cut into horizontal strips and probed with the relevant primary and secondary antibodies and immunoreactivity was detected by ECL.

### Effect of exogenous heparin on FGF4-dependent phosphorylation of FRS2α, MAPK1 and MAPK3

It is conceivable that the interaction of FGF4 with HS in the pericellular matrix of Rama-27 cells and subsequent diffusion of the FGF away from the such HS to the FGFR could be responsible for some observed concentration- and time-dependent changes in phosphorylation of FRS2α at Y436, Y196, MAPK1 and MAPK3. Exogenous heparin will compete for binding to HS and is perfectly competent to enable the assembly of a ternary signalling complex of FGF ligand, FGFR and HS[9, 10, 27]. To determine whether the interaction of FGF4 with HS in the pericellular matrix of Rama 27 fibroblasts influenced the concentration-dependence of downstream phosphorylation of FRS2α, MAPK1 and MAPK3 we included 1 µg/mL heparin. Quiescent Rama 27 fibroblasts were stimulated with FGF4 in the presence of 1 µg/mL heparin. Under these conditions the heparin will outcompete endogenous cell HS and any effects of FGF4 interacting with the latter will be markedly reduced, if not abolished. Heparin itself had no detectable effect on the phosphorylation of FRS2α, although as previously noted it caused an increase in phosphorylation of MAPK1 and MAPK3 (Fig. 5)[9, 26, 27, 29]. Phosphorylation of FRS2α, MAPK1 and MAPK3 stimulated by 10 pg/mLFGF4 in the presence of heparin was not detectable after 15 minutes and 60 minutes (Fig. 5), consistent with the result obtained in the absence of heparin (Figs 2, 3A, B). The phosphorylation of FRS2α, MAPK1 and MAPK3 at 15 min in the presence of heparin was similar to that in its absence Figs 3C, D and 5). However, at 60 min, heparin did appear to reduce to some extent the phosphorylation of FRS2α at Y196 and of MAPK1 and MAPK3 (Figs 3C, D and 5). With FGF4 at 10 ng/mL it appeared that heparin again attenuated the phosphorylation of FRS2α and MAPK1 and MAP3 observed at 60 min, relative to that at 15 min (Figs 3E, F and 5). At 100 ng/mL FGF4, heparin appeared to dampen somewhat the phosphorylation of FRS2α at 60 min relative to that seen at 15 min (Figs 4 and 5). These data suggest that there may be an influence of the interaction of FGF4 with pericellular matrix and the dose-and time-dependence of the phosphorylation of FRS2α and MAPK1and MAPK3.

**Figure 5:**
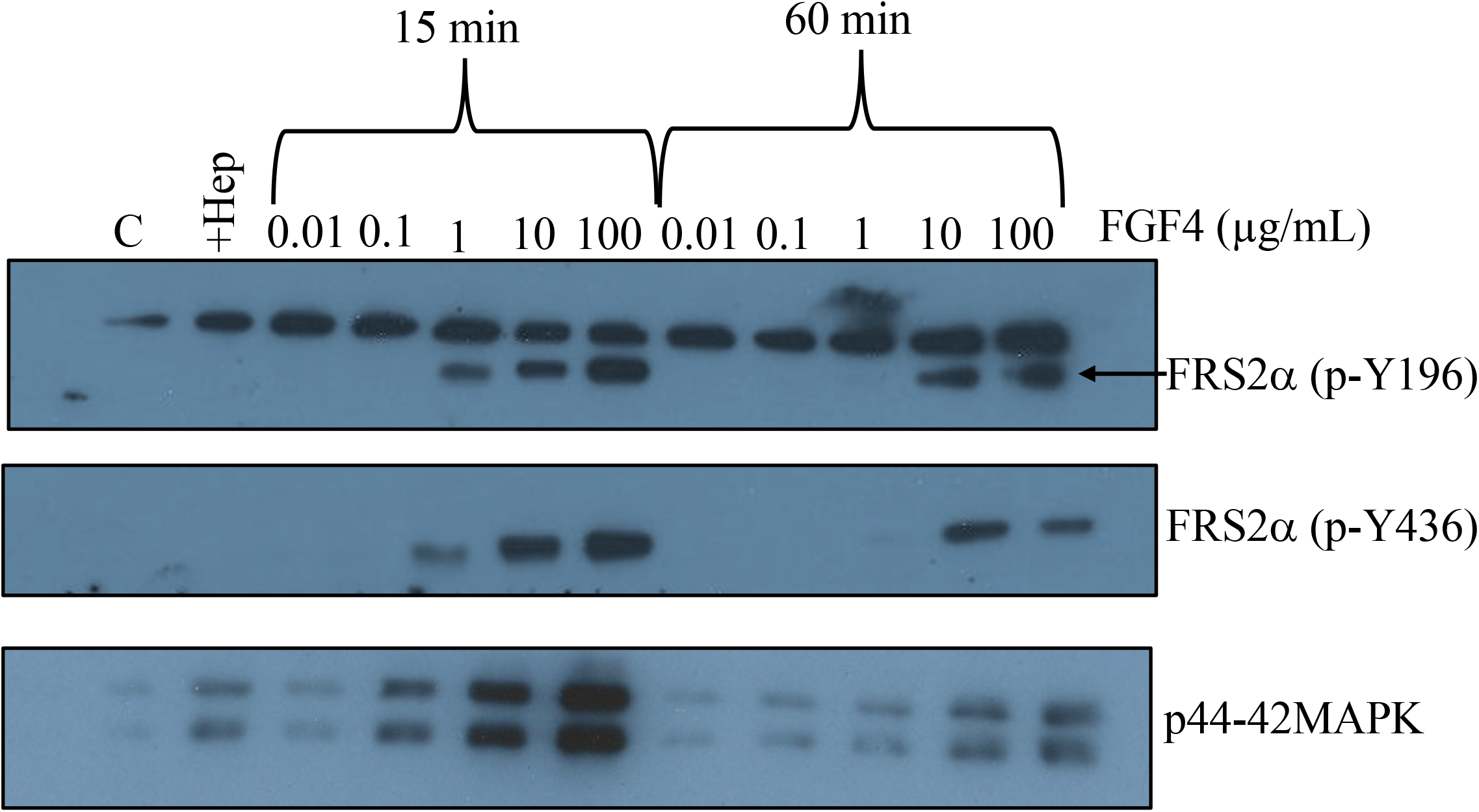
The stimulation of phosphorylation of FRS2α (p-Y436, p-Y196), MAPK1 and MAPK3 by FGF4 in the presence of heparin. Quiescent Rama 27 fibroblasts were incubated with the indicated concentrations of FGF4 and 1 µg/mL heparin. Cells were collected by scrapping in sample buffer after 15 min and 60 min and polypeptides separated by SDS-PAGE. Strips of the nitrocellulose membrane were cut after electro transfer and probed with the relevant antibodies and immunoreactivity identified by ECL

## Discussion

In common with all paracrine FGFs, FGF4 binds to cellular HS and the signalling complex that is responsible for stimulating cell division comprises FGF ligand, FGFR and HS. The pivotal role of FGF4 gradients in vertebrate limb development illustrate its importance in the formation of body structures. As noted, however, there is a paucity of information concerning the concentration- and time-dependence of intracellular signals relating to cell division that are stimulated by FGF4. The Rama 27 fibroblasts used here only possess detectable FGFR1, which allows a comparison with the substantial data accumulated with these cells on FGF2. FGF4, like FGF2, elicited phosphorylation of F`RS2α, MAPK1 and MAPK3. It also induced the phosphorylation of AKT, which had not been examined previously in these cells. FGF4 was less potent than FGF2, in that the early phosphorylation of FRS2α, MAPK1 and MAPK3 reached a maximum at ∼ 10 ng/mL FGF4 and is half maximal at ∼ 1 ng/mL (Fig. 3), whereas 0.3 ng/mL FGF2 elicited maximal phosphorylation of these proteins after 15 min [9, 27]. In contrast, FGF2 and FGF4 were shown to be equipotent in terms of their mitogenicity in Baf cells transfected with a specific FGFR isoform[30]. The differences may arise from the assays (phosphorylation *versus* mitogenicity), though for FGF2 in Rama 27 fibroblasts these two parameters had very similar dose responses[9]. Alternatively, the cell systems themselves may underlie the differences in potency. Rama 27 fibroblasts are not transfected and express what might be considered a ‘natural’ complement of FGFR1 (of the order of 1000s per cell) and HS-containing pericellular matrix [29, 31, 32] whereas the Baf3 cells are necessarily transfected (they do not express FGFRs) and exogenous heparin has to be present since they also lack HS [30]. Moreover, ∼100-fold higher concentrations of heparin are required in the latter cells than in Rama 27 cells that have been rendered devoid of sulfated HS [9, 26, 27] [30] Therefore, the differences in activity of FGF2 [9] and FGF4 (Fig. 3) observed with Rama 27 cells may be a closer reflection of the relative potency of the two FGFs *in vivo*.

Interestingly, oscillations of the phosphorylation of FRS2α, MAPK1 and MAPK3 elicited by FGF4 were observed. Since these were at early times in quiescent cells, the underlying mechanism will not be due to cell division, as observed in zebrafish and *Drosophila[33]*. However, the present data suggest that the activation of at least some signalling pathways by FGF4 may be temporally dynamic, which could thus play an important role in the regulation of development by this growth factor. There was evidence for a biphasic response of Rama 27 fibroblasts to FGF4 (Figs 3 and 4), but it was much less marked than reported for FGF2[20] [9, 20-22] . Moreover, the concentrations of FGF4 at which the phosphorylations of FRS2α, MAPK1 and MAPK3 were attenuated were more than 10-fold higher than seen for FGF2 and it remains to be determined whether such concentrations are attained *in vivo* for example in the vicinity of a cell(s) acting as the source of FGF4.

HS in the pericellular matrix binds a high proportion of exogenously added FGFs, such that the concentration added to the cell culture medium is notional, in that the local concentration of the FGF in pericellular matrix will be higher. However, this again will be a notional concentration, since the FGF is largely associated with HS rather than free in solution[4, 31, 32] FGFs bound to HS in the pericellular matrix are effectively trapped, but readily diffuse[4] [31] by mechanisms that may involve sliding and hopping (as discussed in [34, 35]). The concentration of exogenous heparin used here would compete for binding FGF4. Thus, the signalling complex will consist of FGFR1, heparin and FGF4 and there will little if any effect of pericellular matrix HS on the FGF4. In particular, since FGF4 will not be bound to pericellular matrix HS, its engagement with FGFR1 will not depend on its diffusion within pericellular matrix. Interestingly, the presence of heparin reduced the potency of FGF4 measured by the phosphorylation of FRS2α, MAPK1 and MAPK3. This may be a consequence of a difference in the mode of formation of the complex, a soluble FGF4: heparin complex binding FGFR1 compared to FGF4 diffusing between HS binding sites in pericellular matrix to engage FGFR. Alternatively, this may be an effect of concentration: the effective concentration of FGF4 is unrelated to its solution concentration, since the FGF4 is normally in a heterogeneous system, HS bound and transiently in solution, which is also very crowded.

## Acknowledgements

We acknowledge financial support from the University of Tabuk, Saudi Arabia (MAS), EC FET-OPEN programme ArrestAD (EAY and DGF).

## Supplementary information

**Figure S1.**
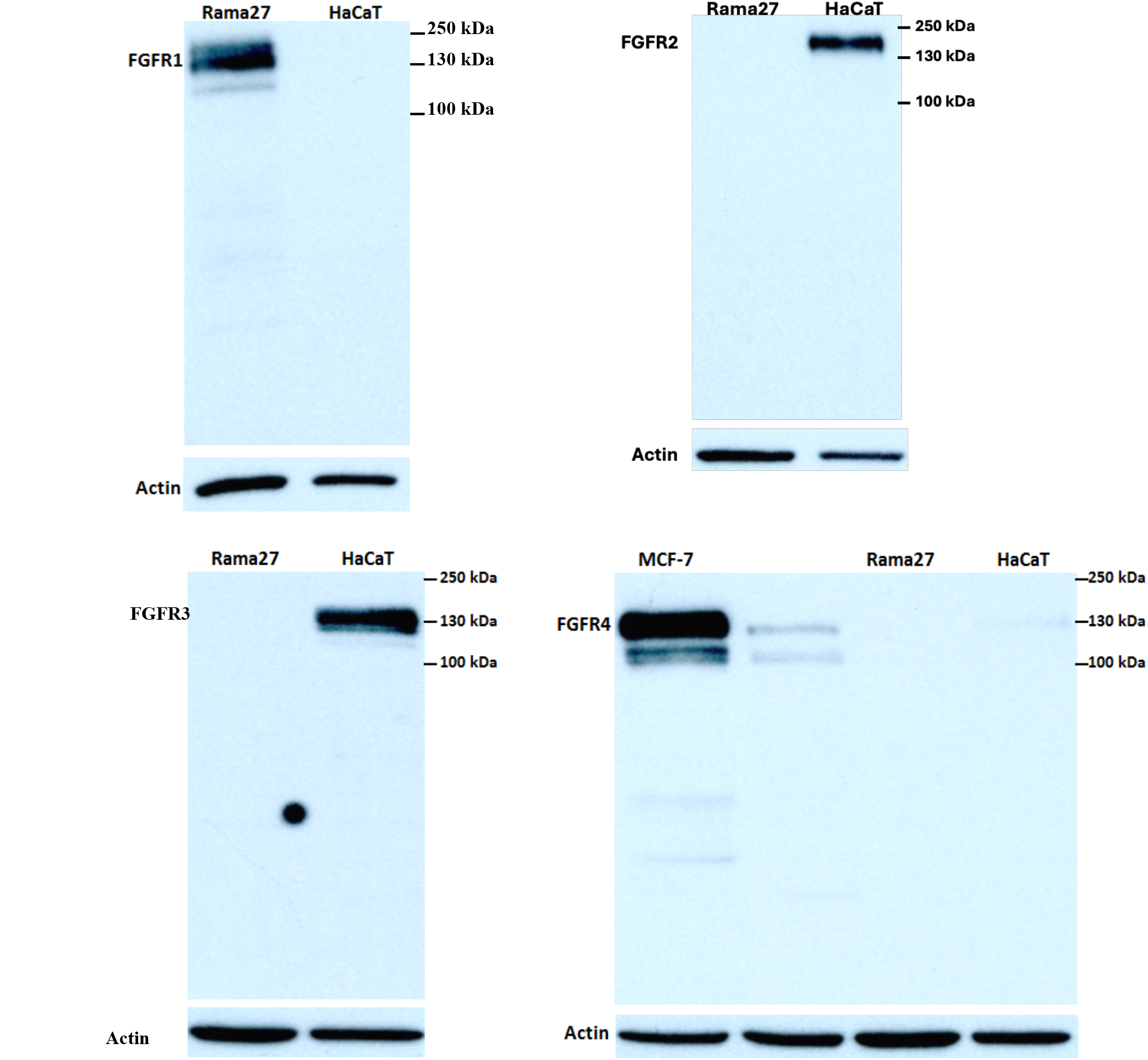
Western blots of FGFRs in Rama 27 fibroblasts. The FGFR is indicated in the figure panels. FGFR1 is the full western blot corresponding to Figure 1. The human keratinocyte cell line, HaCaT was used as a positive control for FGFR2 and FGFR3 and the human mammary carcinoma cell line MCF as a positive control for FGFR4

